# Emergence and spread of the barley net blotch pathogen coincided with crop domestication and cultivation history

**DOI:** 10.1101/2023.07.28.550921

**Authors:** Demetris Taliadoros, Alice Feurtey, Nathan Wyatt, Pierre Gladieux, Timothy Friesen, Eva Stukenbrock

## Abstract

Fungal pathogens cause devastating disease in crops. Understanding the evolutionary origin of pathogens is essential to the prediction of future disease emergence and the potential of pathogens to disperse. The fungus *Pyrenophora teres* f. *teres* causes net form net blotch (NFNB), an economically significant disease of barley. In this study, we have used 104 *P. teres* f. *teres* genomes from four continents to explore the population structure and demographic history of the fungal pathogen. We showed that *P. teres* f. *teres* is structured into populations that tend to be geographically restricted to different regions. Using Multiple Sequentially Markovian Coalescent and machine learning approaches we demonstrated that the demographic history of the pathogen correlates with the history of barley, highlighting the importance of human migration and trade in spreading the pathogen. Exploring signatures of natural selection, we identified several population-specific selective sweeps that colocalized with genomic regions enriched in putative virulence genes, and loci previously identified as determinants of virulence specificities by quantitative trait locus analyses. This reflects rapid adaptation to local hosts and environmental conditions of *P. teres* f. *teres* as it spread with barley. Our research highlights how human activities can contribute to the spread of pathogens that significantly impact the productivity of field crops.

## Introduction

Fungi cause devastating diseases in crop plants and can be dispersed across continents by agricultural trade (1,2). Understanding the evolutionary history of fungal pathogens and the mechanisms underlying their emergence and spread is essential in preventing future epidemics in agroecosystems. Notably, information on historic and current evolutionary trajectories can be key in the development of regulation for re-engineer crops and agroecosystem to improve epidemiological surveillance and prevent potential outbreaks (3).

*Pyrenophora teres* f. *teres* is a widespread fungal pathogen of barley causing the disease “net form net blotch” (NFNB). Infection of susceptible barley leaves occurs by the production of specialized infection structures, appressoria, whereby the fungus penetrates the cuticle and cell wall of epidermal cells. The characteristic net blotch symptoms arise through necrosis which is induced rapidly after initial pathogen invasion of the host leaf (4). Net form net blotch disease occurs in all barley-producing regions of the world, and has been reported in several African countries (5–7), West and East Asia (8,9), Europe (10), North and South America (11,12), and Australia (13). The pathogen has been known to humans for centuries, initially described as *Helminthosporium teres* (Sacc.), however insights into the population biology and demographic history of *P. teres* f. *teres* are scarce.

Barley was domesticated in the Fertile Crescent approximately 10,000 years ago and was later introduced to North Africa and Eurasia by Neolithic farmers (14). North Africa has been proposed as a center of diversity of wild barley (15–18). In the past few centuries, barley has further been dispersed with European migrants to the Americas, Australia, and South Africa (19). Altogether, the dispersal history of barley is associated with human activities and reflects the spread of cereal cultivation through historical and modern trading routes.

Plant domestication has been associated with the emergence of new fungal pathogens (20). Population genetic and evolutionary studies have been used to track down the origin and dispersal history of important crop pathogens. The fungal wheat pathogen *Zymoseptoria tritici* emerged at the onset of wheat domestication and was dispersed with wheat farming during the Neolithic and much later with European migrants (21,22). Likewise, the center of origin of the maize infecting smut fungus *Ustilago maydis* likely also coincide with the center of maize domestication in Central and South America (23). Several other examples underline the importance of domestication and agricultural trade in shaping the evolution and dispersal of crop pathogens (24–26). In modern times, breeding of new crop species has also driven the rapid evolution of new pathogen species, such as mildew pathogens affecting the hybrid crop triticale (27).

The dispersal of pathogens across continents may be accompanied by local adaptation to distinct environmental conditions and/or to specific management practices such as local crop varieties or, in more recent times, fungicides (22,28). Signatures of adaptation can be identified in population genomic data as “selective sweeps”, which are genomic regions with low genetic variation and elevated linkage disequilibrium (29,30). Several statistical methods have been developed to distinguish signatures of selective sweeps from other scenarios that can influence patterns of variation along genomes, such as demography, recombination rate variation, and population structure (30). Prime candidates for signatures of strong and recent positive selection in plant pathogens are genes that encode effector proteins (31–34), which are small proteins secreted by pathogens to manipulate their host’s physiology. Effector proteins play determining roles in the suppression of plant immune responses and have been a major focus in molecular plant pathology research (35). Effector genes can be predicted from genome sequences as they typically encode a signal peptide targeting them for secretion into the plant apoplast or translocation into the host cytoplasm. Moreover, effectors are typically cysteine-rich and specifically expressed during host invasion (36). Several effector genes have been identified by quantitative trait locus analysis (QTL) or genome-wide-association studies (GWAS) underlining the importance of genome data in the discovery of virulence mechanisms (37).

In the present study, we analyzed the haploid genomes of 104 *P. teres* f. *teres* isolates derived from different barley fields worldwide. Based on single nucleotide polymorphisms (SNPs), we characterized the population structure and inferred the demographic history of the pathogen. We specifically investigated patterns of early lineage divergence using different methods for demographic inference. Our analyses provide strong evidence for a recent origin and dispersal of *P. teres* f. *teres*, likely coinciding with the domestication and dispersal of barley by Neolithic farmers. We, moreover, investigated signatures of recent natural selection and found an overlap between signatures of selective sweeps and putative virulence factors previously identified by quantitative trait locus (QTL) analyses (38,39). Our study reveals the recent emergence of an important crop pathogen along the domestication and dispersal of its host and underlines the impact of human activities and agriculture on the evolution and spread of new diseases.

## Results

### Generation of a population genomic dataset of *P. teres* f. *teres*

To study the geographical population genetic structure of *P. teres* f. *teres* and to infer the recent history of this emerging barley pathogen, we generated a population genomic dataset comprising sequence data of 104 isolates from cultivated barley in six countries across four continents (Africa, America, Central Asia, and Europe) (Table S1). The genomes were sequenced with Illumina technology and sequencing reads were mapped to the reference genome of *P. teres* f. *teres* (40) to identify SNPs. The average read coverage of genomes was 21X, and we identified a total of 1,092,635 high-quality SNPs among the 104 isolates. Further summary statistics related to the read mapping and variant calling are summarized in Table S2.

### Phylogenetic relationship of *Pyrenophora* species from barley and other grass hosts

Because barley can be infected by other closely related *Pyrenophora* species, including *P. teres* f. *maculata*, we first reconstructed the phylogenetic relationships between the isolates included in our study and other barley infecting *Pyrenophora* species to ensure the species identity of our isolates. Our analyses included a set of isolates collected from wild barley exhibiting spot lesions in the Monterey Peninsula (California), which allowed us to compare genetic diversity in *P. teres* f. *teres* populations of cultivated and wild barley.

Using sequence information from four gene loci (ITS, LSU, *tub2*, and *tef1-a*) (41,42), we found that the Californian population from wild barley represents a separate lineage or species of *Pyrenophora* that is more closely related to *P. graminea* than *P. teres* (Figure 1A). The Californian *Pyrenophora* population thereby provided us with an ideal outgroup for further analyses of the *P. teres* f. *teres* populations.

**Figure 1:**
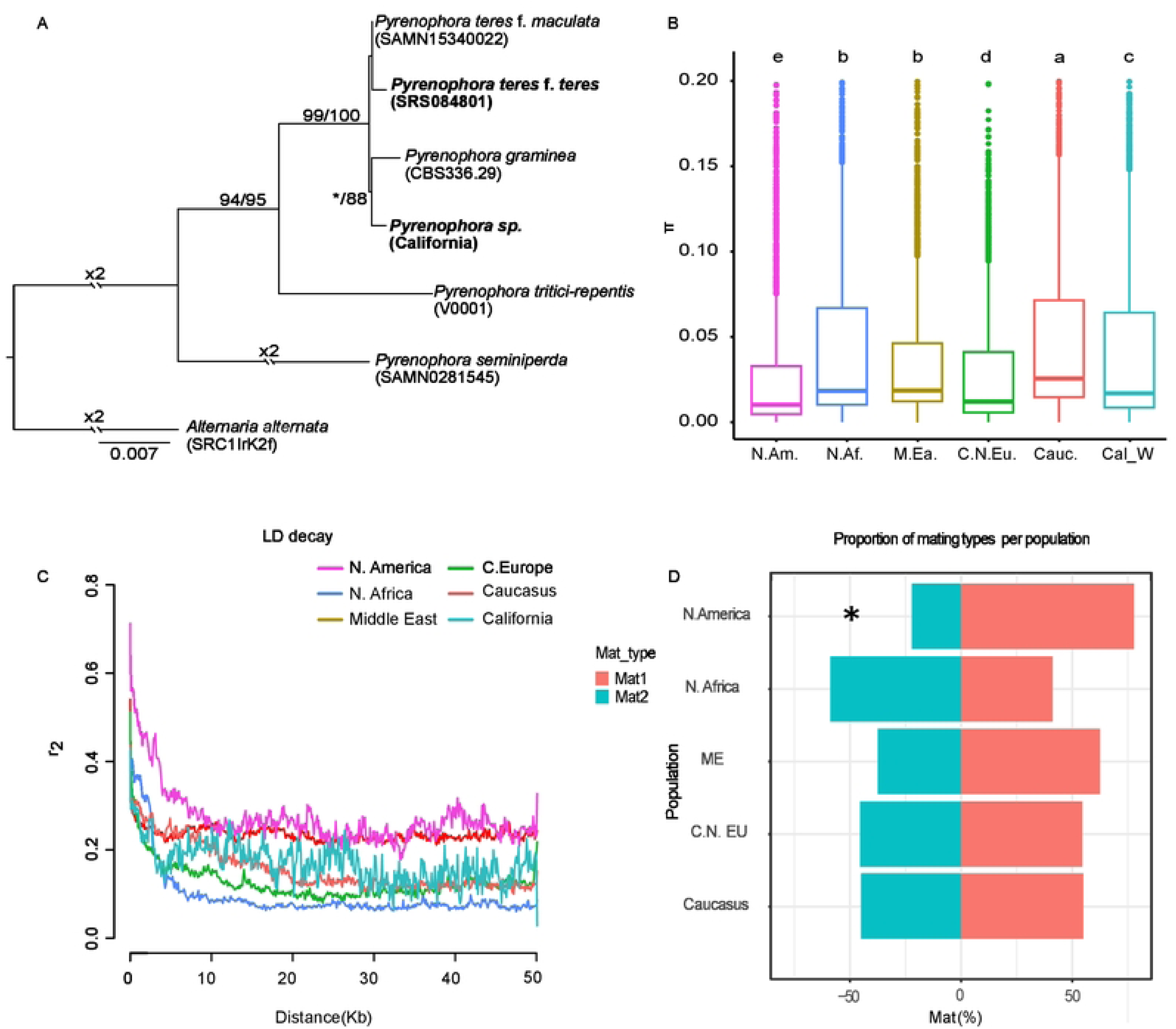
Collection of *Pyrenophora teres* isolates across continents for the inference of pathogen population structure and dispersal. A) Inference of the phylogenetic relationship of closely related *Pyrenophora* species, including isolates of different species originating from barley. The tree was built with nucleotide sequence alignments of the ITS, tub2, LSU, and tef1 regions obtained from five *Pyrenophora* species (Maximum likelihood inference, loglikelihood: −12,351.458). Numbers reflect maximum likelihood and maximum parsimony bootstrap, respectively. *Alternaria alternata* was defined as root. B) Nucleotide diversity of *P. teres* f. *teres* populations in each geographic region. Kruskal-Wallis test with post-hoc pairwise Wilcoxon was used to identify significant differences (p < 0.05) between the groups (Table S3). C) Linkage disequilibrium decay for each population. D) Percentage of the two mating types occurring in each location. Asterisk indicates significant departure from the 1:1 ratio (chi-squared test, p-value: 0.05).

### The Caucasian population of *P. teres* f. *teres* exhibits higher nucleotide diversity

The level of standing genetic variation present in populations can give insight into their demographic history. We compared the nucleotide diversity among the six geographical *P. teres* f. *teres* populations using the Kruskal-Wallis test, which revealed significant differences between the populations (p-value < 2.2e-16). Subsequently, we performed a pairwise Wilcoxon test to assess significant differences between the populations (Table S3). Our analysis showed that the Caucasus population harbours significantly higher genetic diversity (mean π Caucasus = 0.0518) than every other population. Furthermore, even though the Middle Eastern and North African populations showed significantly lower genetic diversity (mean π: 0.0464) that the Caucasian population, they showed significantly higher diversity than the European and the North American populations. The North American population had the lowest level of nucleotide diversity (mean π: 0.0301) (Figure 1B).

Similar to genetic diversity, we used the Kruskal-Wallis test and the pairwise Wilcoxon test (Table S4) to assess differences in Tajimás D values. Indeed, we found that Tajimás D values were significantly different between *P. teres* f. *teres* populations (Kruskal-Wallise test, p-value < 2.2e-16). Our results showed that Tajimás D is close to zero for Caucasus (mean D: 0.04), suggesting a mutation-drift equilibrium (Table 1, Figure S1). Middle East and North American populations showed significantly higher Tajimás D values compared to the other populations (mean D: 0.70 and 0.36, respectively), indicating that these populations have undergone a bottleneck recently. In contrast, the North African population showed the lowest Tajimás D value (−0.27) which may reflect a recent expansion for the North African population. Considering that the Caucasian population exhibits the highest level of diversity with a frequency of genetic variants that reflects a mutation – drift equilibrium, we hypothesize that this population is older than the other populations in our dataset.

**Table 1:** Summary statistics of the genetic clusters of *P. teres* f. *teres*.

| Cluster name | No of individuals | Isolate origin | $\pi$ | $\Theta_w$ | Tajima | r2/2 | Dist r2/2 (Kbp) | Dist r2 < 0.25 (Kbp) | Ne ($\pi$) | Ne ($\Theta_w$) |
| --- | --- | --- | --- | --- | --- | --- | --- | --- | --- | --- |
| Middle East | 7 | Iran | 0,003 | 0,001 | 0.695 | 0,27 | 0,7 | 153 | 33200,000 | 13772,667 |
| North Africa | 23 | Morocco | 0,006 | 0,002 | -0,27 | 0,24 | 2,7 | 151 | 61311,111 | 24257,667 |
| Caucasus | 18 | Azerbaijan (11),<br>Denmark (3),<br>Iran (2),<br>Morocco (2) | 0,005 | 0,002 | 0,04 | 0,26 | 7,3 | 152 | 58366,667 | 23468,889 |
| Central and North Europe | 20 | France(18),<br>Denmark (1),<br>Azerbaijan (1) | 0,005 | 0,002 | -0,10 | 0,22 | 1 | 150 | 55922,222 | 20547,444 |
| North America | 20 | ND | 0,004 | 0,002 | 0,36 | 0,36 | 3,9 | 164 | 46544,444 | 18372,778 |
| [1] $\pi$: Estimator of genetic diversity based on the pairwise nucleotide differences of a genetic region (Nei and Li, 1979) | | | | | | | | | | |
| [2] $\Theta_w$: Waterson's $\Theta$ (Waterson, 1975) | | | | | | | | | | |
| [3] Tajima's D (Tajima, 1989) |  |  |  |  |  |  |  |  |  |  |
| [4] Correlation coefficient |  |  |  |  |  |  |  |  |  |  |
| [5] Ne ($\pi$): $\pi/2\mu^*$ | | | | | | | | | | |
| [6] Ne ($\Theta_w$): $\Theta_w/2\mu$ | | | | | | | | | | |

Varying extent of linkage disequilibrium (LD) among fungal populations can also inform about the frequency of sexual reproduction, and reflect different ages of populations. More recently founded populations, and populations with lower frequencies of sexual reproduction, will typically exhibit a greater extent of LD compared to older or sexually recombining populations. We found considerably longer linkage blocks in the North American population, for which the LD statistic r^2^ was reduced to half of its maximum value at 3.9 Kbp (Figure 1C, Table 1). The long LD blocks observed for the North American population suggest that this population was founded more recently.

*Pyrenophora teres* f. *teres* has a heterothallic mating system implying that mating only occurs between individuals of opposite mating types, Mat1-1 and Mat1-2 (43). To further investigate geographic variation in the frequency of sexual reproduction, we examined the mating type ratio, which is expected to be equal to one under random mating. We used the software SPAdes (44) to *de novo* assemble genomes and thereby validate and compare the frequency of mating type loci. The null hypothesis of random mating could not be rejected for the *P. teres* f. *teres* populations except the North American population, for which we found a significant departure from the expected 1:1 ratio of mating types (Mat1-1:Mat1-2 = 3.5, Chi-squared test, p = 0.0184) (Figure 1D, Table S5). These analyses suggest that that *P. teres* f. *teres* is regularly undergoing sexual reproduction throughout most of its range. The skewed mating type frequency in the In North American population may reflect a more pronounced contribution of asexual reproduction.

### Populations of *P. teres* f. *teres* are geographically structured

We characterized the population genetic structure of *P. teres* f. *teres* based on complementary methods using genome-wide SNP data. Firstly, we investigated the extent of clustering using a principal component analysis (PCA) (Figure 2A). The PCA mostly separated isolates according to their geographical origin. We further explored population structure by generating a Neighbour-net network with SPLITSTREE v. 4 and by inferring the extent of shared ancestry using an ADMIXTURE analysis (45). In the ADMIXTURE analysis, the Cross-Validation error used to select the most appropriate number of clusters (K) was minimized at K = 6 (Figure 2B, Table S6). The genetic clusters inferred from the ADMIXTURE analysis corresponded to five clusters mostly circumscribed to North America, North Africa, the Middle East, Europe, and Caucasus (Figure 2B), and a cluster of three individuals restricted to the Caucasus referred to as the Caucasus-2 cluster. The European cluster, referred to as Europe+, was also present in the Caucasus (one isolate), and the Caucasus cluster, referred to as Caucasus+ was also present in the Middle East, in North Africa, and in Northern Europe. The ADMIXTURE analysis also revealed 15 individuals with shared ancestry in multiple clusters, suggesting admixture. Most of the individuals showing mixed ancestry (9/15) were derived from the Caucasus, where multiple clusters coexist. Furthermore, three isolates with mixed ancestry were derived from Europe, two from North Africa, and one from the Middle East.

**Figure 2:**
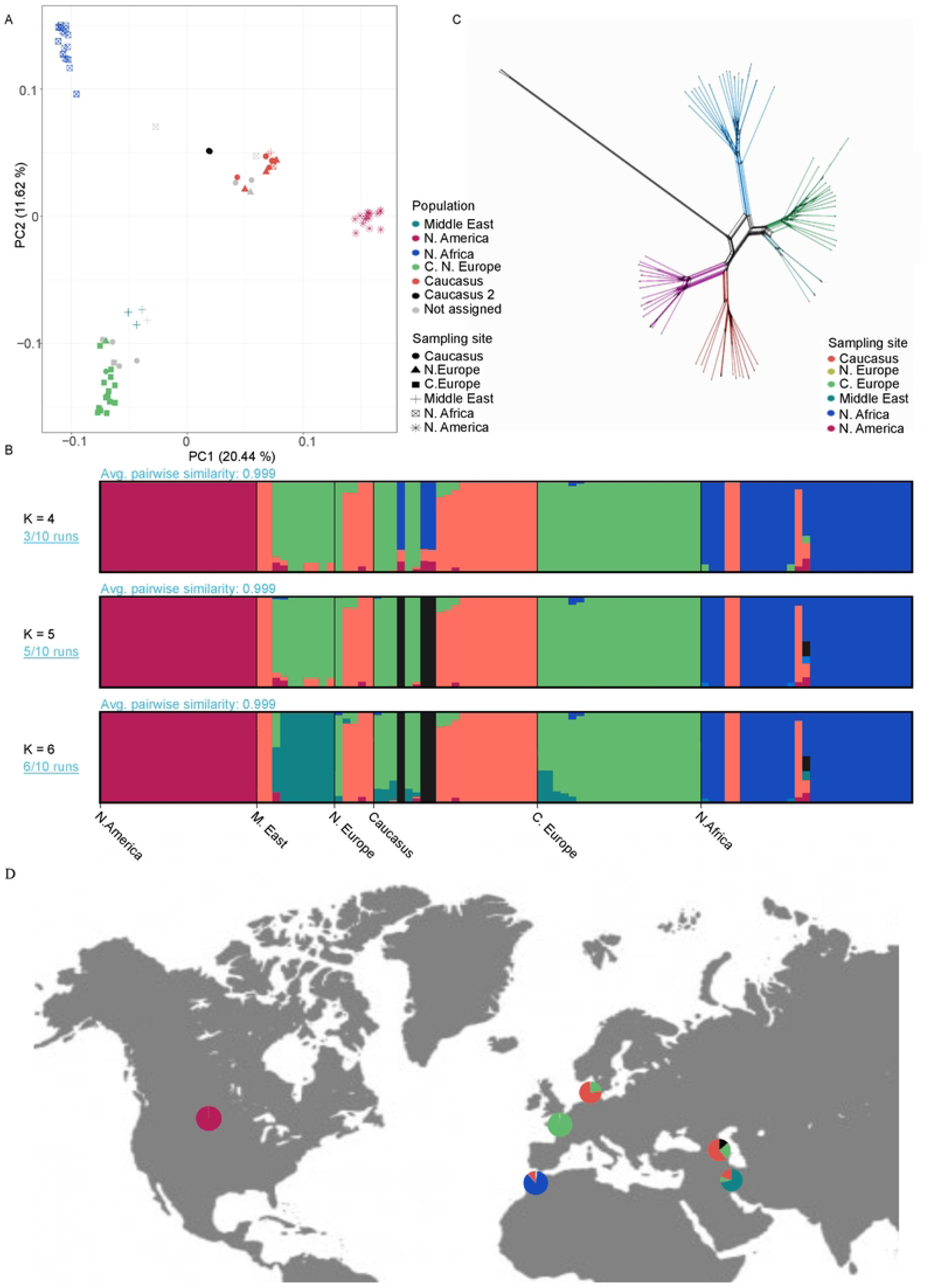
Global population structure of *P. teres* f. *teres.* A) PCA analysis, where shape reflects the origin of the isolate and colour reflects the genetic cluster set with ADMIXTURE at K = 6. B) The program ADMIXTURE was used to compute population structure between the six geographical populations. Most fit number of hypothetical ancestral groups was identified as six based on the cross-validation method (Table S6). Here we present patterns of four, five, and six hypothetical ancestral groups. C) Neighbour-Net tree generated from SNP data from the *P. teres* f. *teres* populations. The branch colour reflects the genetic cluster set with ADMIXTURE at K = 6. D) World map shows the distribution and contribution of the genetic clusters identified by ADMIXTURE at K = 6 at the sampling sites.

The Neighbour-net phylogenetic network essentially revealed the same clusters as the ADMIXTURE analysis. All clusters were connected by reticulations indicating homoplasic mutations caused by incomplete lineage sorting or historical gene flow (Figure 2C). The Caucasus-2 cluster was connected to other lineages by a long, non-reticulated branch, consistent with a relatively long history of isolation from other clusters (Figure S2).

We applied a Mantel test to determine if genetic distance, simply measured as pairwise messmates across the genome, is correlated with geographic distance between the isolates (46), and indeed confirm that geography explains some of the variation between clusters as spatial and genetic distances are correlated (Figure S3).

### Phylogenomic analysis suggests an ancient split of the North African *P. teres* f. *teres* population and a Caucasian origin of the North American population

The Middle East, Caucasus, and North Africa are the regions that have the longest history of barley cultivation (16). Domesticated barley was introduced later to Europe and then to America. To test the hypothesis that early dispersal of *P. teres* f. *teres* occurred simultaneously with the spread of barley cultivation we inferred the evolutionary relationships between the populations of the pathogen. To this end, we have constructed a population tree, using polymorphism-aware models in IQ-TREE, using the Californian population as root for the *P. teres* f. *teres* populations (Figure 3). In this analysis, based on the full complement of polymorphisms, we found two major population splits: one lineage comprising the North African, Middle Eastern, and European population and another lineage comprising the Caucasus and North American populations. Within the former lineage, the branching harbouring the North African population diverged earlier than the branches harbouring the Middle Eastern and European populations. The clustering of Caucasus and North American populations suggested a Caucasian origin of the North American population.

**Figure 3:**
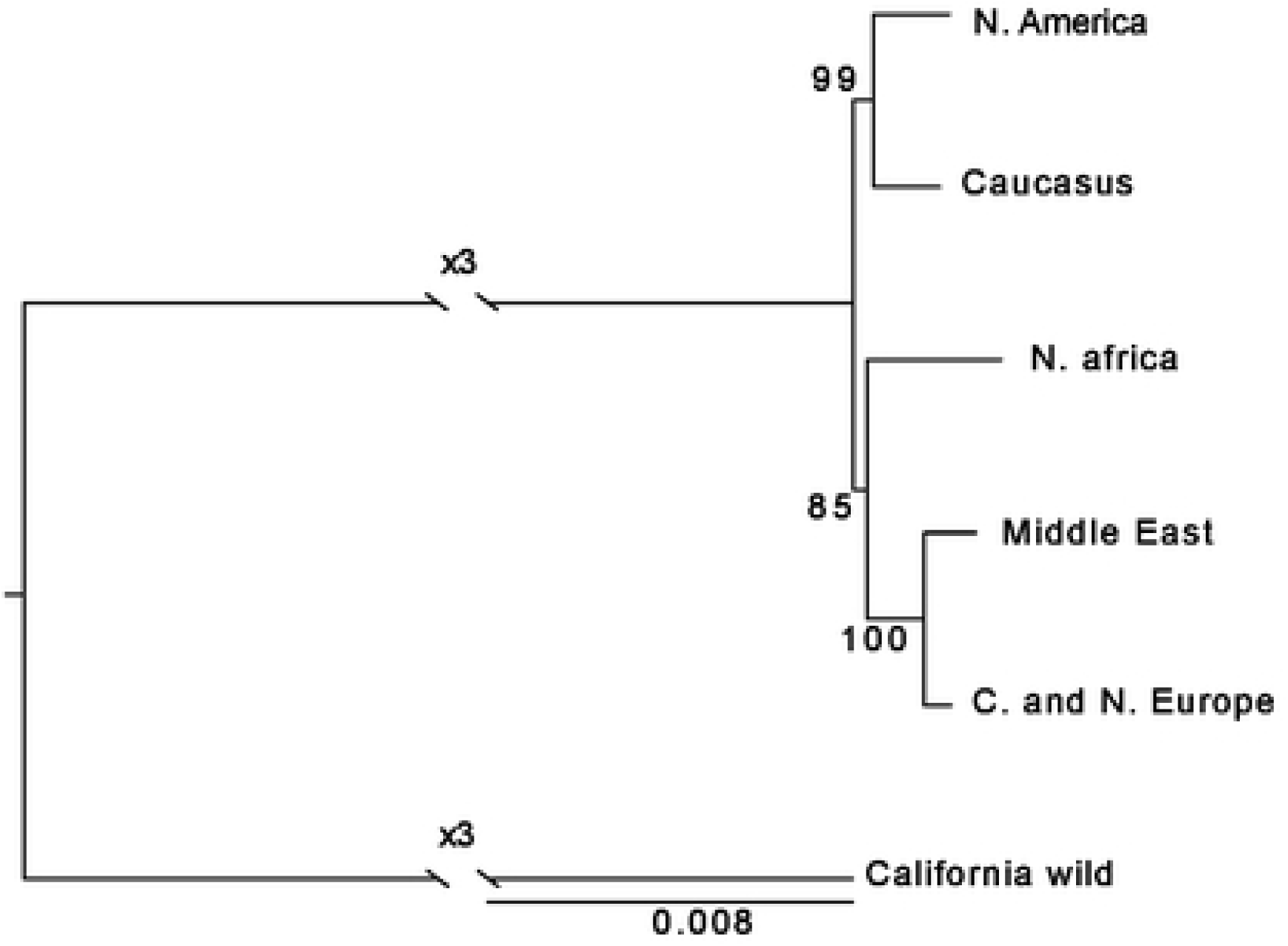
Evolutionary relationship between *P. teres* f. *teres* populations. A) Phylogenetic tree using polymorphism-aware models (PoMo) (Maximum likelihood inference, loglikelihood: −857128.545) to assess the evolutionary relationship between *P. teres* f. *teres* populations. Branch numbers reflect maximum likelihood bootstrap values. The tree was rooted using the Californian population. The scale bar represents the expected number of substitutions per site.

### The origin and dispersal of *P. teres* f. *teres* correlates with the early history of barley cultivation

We further investigated the demographic history of *P. teres* f. *teres* populations using Approximate Bayesian computations (ABC) with supervised machine learning implemented in the DIYABC Random Forest software (DIYABC-RF) (47). As the ABC framework requires populations not connected by continuous geneflow, we excluded isolates with shared ancestry in multiple clusters and considered five, non-admixed geographic populations for the analysis. Non-Caucasian and non-European isolates were excluded from the Caucasus+ and European clusters. We compared invasion scenarios in which the origin of each derived population was associated with a demographic bottleneck (see Materials and Methods) (48,49). For each scenario, we assessed the compatibility of the simulated datasets with the observed data using linear discriminant analysis (LDA), by simultaneously projecting simulated and observed data on the first two LDA axes. The overlap between the simulated datasets and the observed data indicated the compatibility of the simulated scenarios and the observed data (Figure S4, S5, S6).

Demographic inference with ABC was performed in three consecutive steps, each step corresponding to a different family of invasion scenarios. For each family of scenarios, LDA confirmed that the chosen conditions were suitable for the random forest analysis. To select the most probable hypothetical scenario from each family, we used a random forest classifier with 1,000 trees. Detailed results of DIY-ABC analyses are provided in Table S7 and Figure S4-S6.

#### Scenarios family 1: Early divergence of Middle East, Caucasus, North African populations

To elucidate the most ancestral splits, we tested a total of 49 invasion scenarios with different ancestries and branching orders among populations from the Middle East, Caucasus and North Africa, regions where barley barley was first cultivated. Nine distinct categories of scenarios with similar topologies were considered. The most probable category of scenarios was “group 9” and scenario 45 (posterior probability of 0.778 and 0.401, respectively) (Table S8). Scenario 45 modeled an initial divergence of *P. teres* f. *teres* populations from the Middle East and the Caucasus, and a subsequent emergence of the North African population from the Middle Eastern population (Figure 4A).

**Figure 4:**
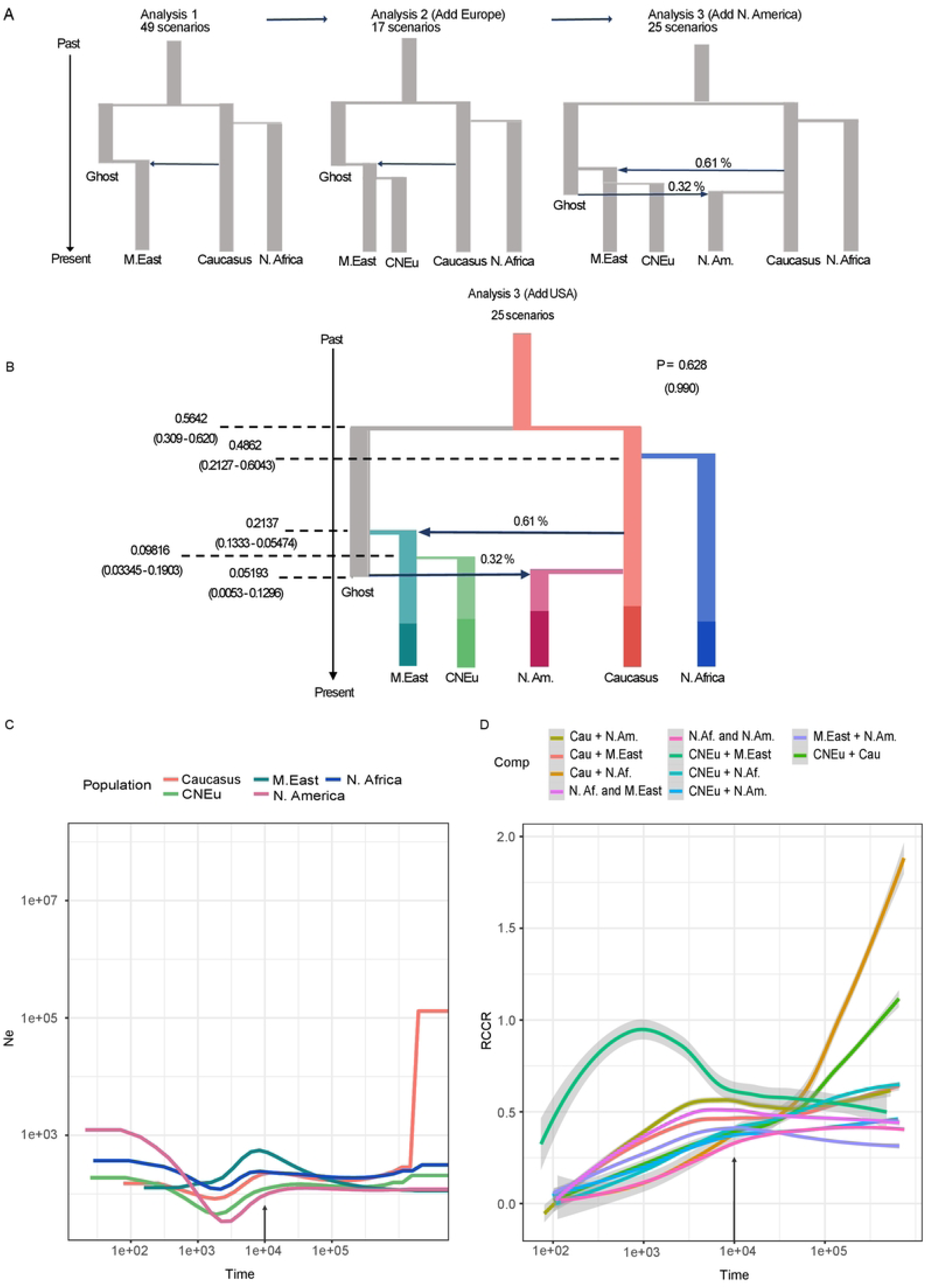
Inference of the demographic history of *P. teres* f. *teres* populations shows an ancient split coinciding with barley domestication and early migration. A) Development of speciation scenarios across three analyses steps and using approximate Bayesian computation and Random Forest analyses implemented in DIY ABC-RF version 1.0. B) The most probable hypothetical evolutionary scenario of the migration routes of *P. teres* f. *teres* in the Middle East, North Africa, Europe, and North America based on the results of three sequential DIY ABC-RF analyses. The parameter “P” indicates the posterior probability of the most probable scenario. In paratheses are shown the probabilities of the different group containing the most probable demographic scenario (see Methods). We considered a classic invasion scenario for the topology building where each derived population passes through a bottleneck as it gives rise to a new population. Predicted time over the current Ne of the most ancestral population (N_Azb_) values inferred with random forest are shown. The 90% CI values for each parameter is provided in parentheses. C) Changes in effective population size for all *P. teres* f. *teres* populations were estimated with MSMC2. The axes were scaled with a mutation rate of 4.5 × 10^−7^ per site per generation, and one generation per year. We have indicated the estimated time of barley domestication with an arrow. D) Relative cross-coalescence rate for all pairs of populations. Five runs of seven randomly selected individuals per population per run were performed. Shown here, we present the trend line fit between the five runs. Gray area around the line indicates the 95% confidence interval. The arrow indicates the estimated time of barley domestication.

#### Scenarios family 2: Founding of the European population through migration from the Middle East

Having determined the branching order among populations from areas of more ancient barley cultivation, we proceeded to examine the more recent history of *P. teres* f. *teres* populations through a second family of scenarios. We compared 17 evolutionary scenarios modeling the relationships among the Central and Northern European populations, and other populations. In line with the population tree presented above, the most probable scenario category was “group 2” and scenario 2 (posterior probability of 0.701 and 0.412, respectively), suggesting that the Central and North European population derived from the Middle Eastern population (Figure 4A).

#### Scenarios family 3: Origin of the North American population

The third family of scenarios modeled the origin of the North American population. According to Scenario 9 from group 5, which had the highest posterior probabilities (0.628 and 0.990, respectively), the North American population was established through admixture between the Caucasus population and an unknown “ghost” population (Figure 4A). This analysis also revealed that the earliest divergence was between the Caucasus and ghost population, which suggest that the Caucasus population is the oldest, and that the Middle East population emerged following admixture between the Caucasus and ghost populations.

#### Parameter inference analysis with DIY-ABC

To estimate the demographic parameters of the scenarios with highest posterior probabilities, we used a random forest with 1000 trees. Time estimates were estimated as the ratio of time over the current effective population size of the predicted most ancestral population, Caucasus population (N_Az_) (47). Our maximum posterior probability estimate of the divergence time between the Caucasus and unsampled ghost populations was 0.5640 generations/N_Az_ (credibility interval [CI] 0.3090 – 0.6200). The North African population split from the Caucasus population 0.4862 generations/ N_Az_ (CI 0.2127 – 0.6043), and the Middle East population emerged by the admixture of the Caucasus population and the ghost population around 0.2137 generations/ N_Az_ (CI 0.1333 – 0.5474). The Central and Northern European population arose from the Middle Eastern population 0.0982 generations/ N_Az_ (CI 0.0334 – 0.1903), and the North American population emerged 0.05193 generations/ N_Az_ (CI 0.0053 – 0.1296) ago (Table 2, Figure 4B).

**Table 2:** Results for DIYABC-RF estimation of population divergence times normalized over the most ancestral sampled population N5 under scenario 9 detailed in Figure 4.

|  |  |  |  |  |  |
| --- | --- | --- | --- | --- | --- |
| **Parameter** | **Prediction\*** | **90% CI** |  | **Global (prior) NMAE** | **Local (posterior) NMAE** |
| t1/N5 | 0,052 | 0,005 | 0,130 | 0,437 | 0,316 |
| t2/N5 | 0,098 | 0,033 | 0,190 | 0,440 | 0,311 |
| t3/N5 | 0,214 | 0,133 | 0,547 | 0,320 | 0,275 |
| t4/N5 | 0,486 | 0,213 | 0,604 | 0,245 | 0,210 |
| t5/N5 | 0,564 | 0,310 | 0,620 | 0,200 | 0,190 |
| Note | *RF analysis included 20,000 simulated data sets and the number of trees was set to 1000. Global (prior) and local (posterior) error rates were estimated using out-of-bag estimators from a sample of 10,000 data randomly chosen in a training set. CI, credibility interval; NMAE, normalized mean absolute error. |  |  |  |  |

### A severe demographic bottleneck in the history of the crop pathogen coincided with the domestication of barley

To further investigate population size variation through time and population split, we applied a Multiple Sequentially Markovian Coalescent (MSMC2) approach (50). Inferences of population size changes and splitting times were performed using all available individuals (Figure 4C). We performed the MSMC2 analysis with five independent runs of 14 randomly selected isolates per population pair (seven isolates per population) (Figure 4D). We found that the effective population size of the Caucasus population was initially the largest, but that the population subsequently experienced a demographic bottleneck. In agreement with the ABC analyses, we also found evidence for an early divergence between Caucasus and the North African population. Considering a mutation rate (μ) of 5.7 × 10^−7^ per base pair (51) and assuming here on average one sexual generation per year (52), these events coincide with the domestication of barley about 7,267 to 14,545 years ago (Table S9).

We computed the Relative Cross Coalescence Rate (RCCR) for pairwise combinations of populations to estimate splitting times (Figure 4D). These analyses provided further support for the close evolutionary relationship between Central Europe and Middle East populations, as also observed in the PCA and NeighborNet tree analyses. Furthermore, the RCCR of the North America and Caucasus populations decays slower, which indicates extensive amounts of geneflow after divergence of the populations. The geneflow and late split between North America and Caucasus populations interfered with the RCCR, is in line with the emergence of the North American population from the Caucasian population as inferred by the ABC analysis. We want to underline that our inference of actual coalescence times is based on assumptions that we considered reasonable for some unknown parameters. For example, the number of sexual cycles of *P. teres* f. *teres* is not known, and might even have varied throughout evolutionary times and host shift events. Nevertheless, the relative estimates of population divergence with two independent methods suggest that the Caucasus has been the center of origin of *P. teres* f. *teres*. Moreover, both methods applied here, provide evidence for an early divergence of a pathogen lineage in North Africa. In summary, our inference of pathogen population history suggests a parallel dispersal of the pathogen alongside its host and emphasizes the fundamental importance of early agriculture on pathogen evolution.

### Recent positive selection has shaped genomic regions encoding putative virulence-related genes

To identify genomic regions that may have experienced selective sweeps during the spread of *P. teres* f. *teres*, we used three methods (SweeD, OmegaPlus, and RAisD) which combine information from the site-frequency-spectrum (SFS) and patterns of LD and π along the genome (53–55). We conducted analyses on each population separately to identify population-specific selective sweeps. These selective sweeps may indicate local adaptation in the pathogen populations.

Demography can greatly impact the distribution of genetic variants along the genome and thereby bias inference of selective sweeps (56). We therefore combined the selective sweep analyses with simulations of genetic variation under different demographic scenarios (see Material and Methods). Multiple regions exhibiting signatures of selective sweeps were identified with the three methods (Table S10 – S15). We compared and combined selective sweeps maps of the three methods to get a final list of candidate regions exhibiting signatures of recent positive selection.

Our final list of sweeps includes a total of 109 regions across all *P. teres* f. *teres* populations, with 20 to 27 selective sweeps per population (Figure 5A, D, Table 3). We identified 42 putative effector genes (40) colocalizing with the 109 selective sweep regions, suggesting that genes encoding virulence related traits such as effectors have been most prone to experience recent positive selection. Some selective sweep regions overlapped while others were unique to distinct populations, possibly representing adaptation to different resistance genes in barley or other local environmental conditions. For example, we found one selective sweep region on chromosome 6 (position 2,856,396 −2,964,731) that is present in all *P. teres* f. *teres* populations, except the Middle Eastern population. This region includes a gene encoding a predicted effector, which represents a candidate for future functional studies (Figure 5B, Table 3).

**Figure 5:**
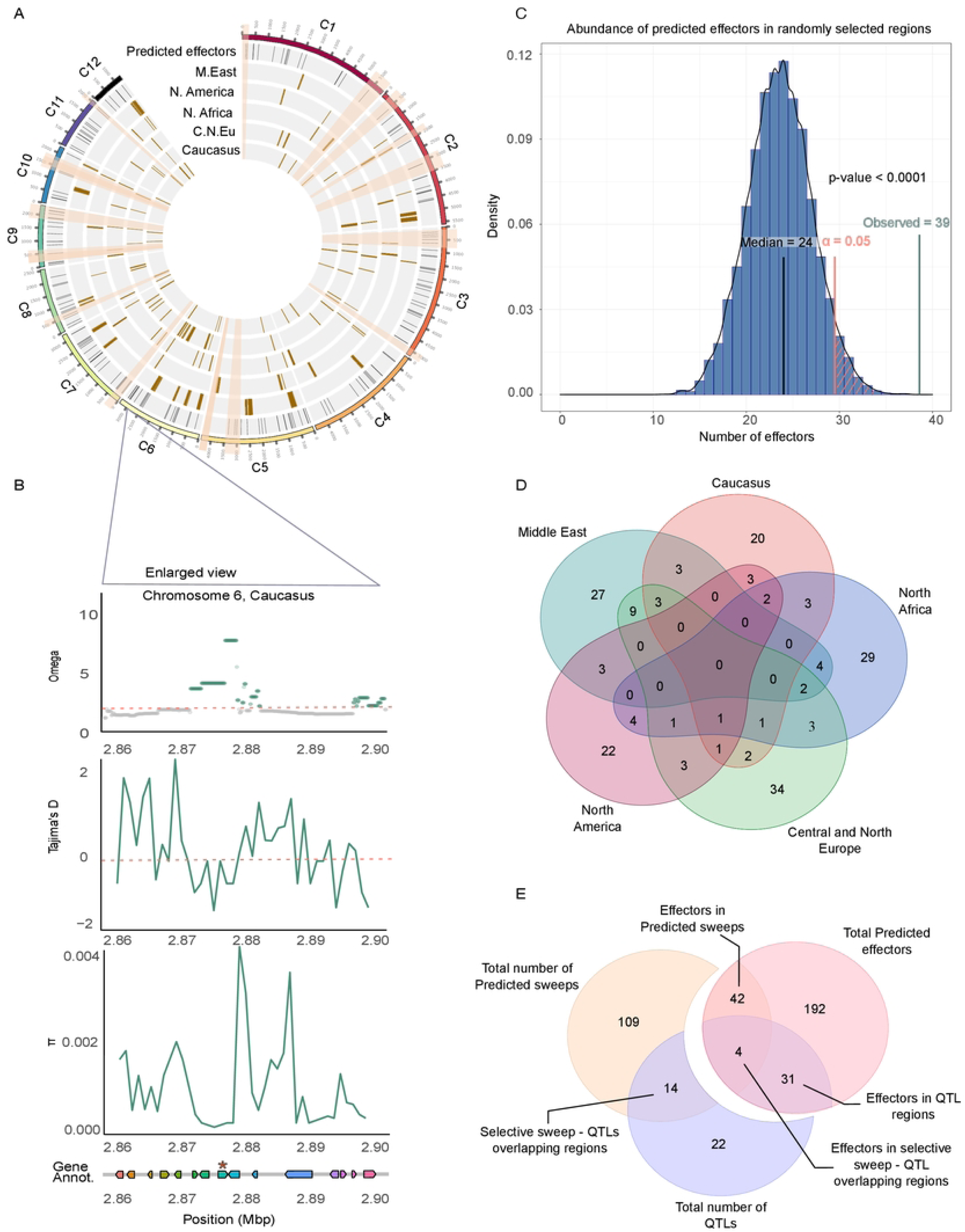
Distribution of selective sweeps across the genome in five *P. teres* f. *teres*. populations. A) Genomic map of selective sweeps for each population. The first track shows coordinates of genes encoding predicted effectors (40). Highlighted are the fourteen QTL regions associated with pathogenicity that were identified in previous studies. (38,39). B) OmegaPlus, Tajima’s D, and nucleotide diversity (π) analyses across a selective sweep region on chromosome 6. Shown is only the Caucasian+ population. This region was identified to be under selection in all populations except the Middle Eastern. At the bottom, the gene and effector annotation presented. A predicted effector situated in the selective sweep region and another effector located in the genomic area. C) To determine if genes encoding putative effectors are enriched in selective sweep regions, we performed an enrichment analysis based on the distribution of predicted effector abundance in randomly selected genomic regions of the same number and length as the selective sweep regions. As many as 10,000 runs of random resampling of genomic regions was perform to validate that effector genes indeed are enriched in regions that have experienced recent positive selection. D) Venn diagram of selective sweep regions shared and unique to the *P. teres* f. *teres* populations. E) Effector content of the selective sweep regions and QTLs. Effector annotation was obtained from Wyatt et al., (2018), and previously reported QTLs associated with virulence in *P. teres* f. *teres* (38,39).

**Table 3:** Summary of total and unique selective sweep regions, as well as, genes and effectors located in the regions per populations. The number of sweep regions, total genes, and effectors only identified in one population are charachterised as unique.

| Population | Total no of Sweep regions | Total no of genes | Total number of predicted effectors | Population-specific sweep regions | Population-specific genes | Population-specific effectors | Selective sweep - QTL overlap | QTLs name | QTL Reference |
| --- | --- | --- | --- | --- | --- | --- | --- | --- | --- |
| Middle east | 27 | 253 | 14 | 11 | 198 | 13 | 1 | VK2 | 40 |
| N. America | 22 | 107 | 9 | 8 | 75 | 8 | 3 | VK1, VK2, AvrHar | 40, 54 |
| N. Africa | 29 | 91 | 5 | 19 | 67 | 3 | 2 | VK2, PttTif1 | 40, 39 |
| C.&N. Europe | 34 | 180 | 5 | 21 | 136 | 5 | 1 | PttPin1 | 39 |
| Caucasus | 20 | 101 | 7 | 11 | 69 | 6 | 2 | VR1, PttCell1 | 40, 39 |
| California (*P. sp*) | 24 | 155 | 5 | 12 | 108 | 4 | 3 | VK1, PttCell1, Pttcell2 | 40, 39 |

We tested if effector genes were significantly enriched in selective sweep regions. To this end, we performed a permutation test to assess the relative abundance of predicted effector genes in the selective sweep regions compared to the rest of the genome. Indeed, we found that the abundance of effector genes in selective sweep regions is higher compared to randomly sampled regions along the genome (Figure 5C).

We furthermore explored previously generated lists of candidate virulence determinants in *P. teres* f. *teres*. Previous studies have used quantitative trait locus (QTL) analyses to identify determinants of virulence on different barley cultivars (38,39). We found that 14 putative virulence related genes (QTL candidates) co-localized with selective sweep regions(38,39). (Figure 5E, Table 3). The QTL candidate region VK2 on chromosome 6 (39) overlapped with a selective sweep region, which was found in each of the populations in the Middle East, North America, and North Africa. The QTL candidate regions VK1 (39) and AvrHar (57) on chromosomes 3 and 5, respectively, co-localized with selective sweep regions predicted in the North American population. Two putative effectors are in the VK1 region on chromosome 3 (Table 3).

Other genes beyond effectors may be important for adaptation of the pathogen to local environmental conditions. We conducted a GO enrichment analysis and found that the selective sweep regions are enriched with genes associated with different GO terms defined for metabolic processes, peptidase activities and protein metabolism among others (Figure S7).

We predict that the selective sweeps in *P. teres* f. *teres* reflect recent adaptation to barley and local agricultural environments. We further addressed divergent adaptation in *Pyrenophora* pathogens on different hosts by comparing selective sweep maps of *P. teres* f. *teres* and the *Pyrenophora* population obtained from wild barley in California. To this end, we considered the windows that showed a composite likelihood ratio (CLR), ω, and μ higher than 99,95 % for significant outliers.

We identified 24 selective sweeps along the genome of the Californian population, including five genes predicted to encode effector genes. Approximately half of the selective sweeps predicted in the Californian population were shared with the domesticated barley-infecting populations suggesting that the same suite of genes is important for virulence on wild and cultivated hosts. This hypothesis is further supported by the fact that some *P. teres* f. *teres* QTL candidates (39) overlap with selective sweep regions in the wild barley pathogen (Table 3). Hereby, also the QTL locus VK1 co-localized with selective sweeps in the wild-barley pathogen.

In summary, the selective sweep analyses identify multiple loci in *P. teres* f. *teres* that have experienced recent positive selection. Functional analyses of candidate genes in these regions may shed light on the adaptation of the pathogen to different barley cultivars.

## Discussion

Understanding the evolutionary origin of crop pathogens is crucial to predict future epidemics. In this study, we addressed the history of the globally occurring pathogen of barley, *P. teres* f. *teres*. We used a global population sample and extensive genome sequencing to assess the population structure and demographic history of the crop pathogen. Our analyses were based on the hypothesis that *P. teres* f. *teres* could have emerged and co-evolved with barley during early crop domestication. Extensive sampling of the pathogen in geographical regions representing the most ancestral history of barley domestication and cultivation (58) allowed us to dissect the early history of *P. teres* f. *teres*. We also characterized the population structure and demography from present-day barley-producing countries, including France, Denmark and the USA. Our detailed population genomic analyses provide evidence for a scenario where *P. teres* f. *teres* emerged in the Middle East at the onset of barley domestication, and subsequently dispersed with Neolithic farmers to North Africa and Europe.

We compared measures of nucleotide diversity among the different *P. teres* f. *teres* populations and observed higher diversity in the Caucasian and North African populations. Notably the North American population represented an overall low nucleotide diversity indicating either a more recent origin of the population or a recent bottleneck.

Next, we applied two independent methods to infer the population histories of *P. teres* f. *teres*. Both methods provide evidence for a scenario where the most ancestral populations of *P. teres* f. *teres* have originated in the Fertile Crescent region. We note that our inferences of population histories based on the Middle East, Caucasus, and North Africa populations, may have been affected by sampling bias as only ten isolates were available from the Middle East, in contrast to 21 and 27 from Caucasus and North Africa, respectively. For the parameter inference of ABC-RF, we have calculated time over the effective population size to reduce the error of parameter inference, as suggested in (47). We computed current effective populations sizes based on the SNP data to rescale parameters with a mutation rate of 4.5 × 10^−7^ (25) and one generation per year (see Materials and methods). Using these values, we estimated that the Caucasian population was founded around 13,000 years ago (CI: 7,267 – 14,546).

The history of *P. teres* f. *teres* not only parallels the evolution of barley. It also parallels the history of a small number of other prominent crop pathogens which have emerged and co-evolved with their host during domestication. Other important pathogens that emerged with their host during domestication include the wheat pathogenic fungus *Zymoseptoria tritici* causing the disease septoria tritici blotch (59), the rice blast fungus *Magnaporthe oryzae* (60), and the corn smut fungus *Ustilago maydis* (61). In these studies, coalescence analyses were used to infer the divergence time between wild and crop-infecting populations of the pathogen and to infer major demographic events, such as bottlenecks that coincide with the domestication of the host. The emergence of *P. teres* f. *teres* was also associated with a considerable population bottleneck probably reflecting strong selection on pathogen individuals with the right gene combination necessary to invade a new host niche.

Interestingly, we find evidence for the early emergence of a distinct *P. teres* f. *teres* population in North Africa. Using the above-mentioned scaling the divergence between pathogen populations in North Africa and Caucasus occurred around 11,400 years ago (CI: 4,991 – 14,181). This scenario is in agreement with the introduction of barley into North Africa by neolithic farmers and the early development of distinct barley varieties (62).

Archaeological remains suggest that barley was cultivated in Central Europe from approximately 6,000 (63) to 4,800 years ago (64). We find evidence that the *P. teres* f. *teres* pathogen accompanied the introduction of barley as our population genomic data suggest the emergence of the European pathogen population occurred around 2,300 years ago (CI: 785 – 4,465). More recently, the North American population has split from all other populations. Our analyses suggest the emergence of the North American population approximately 1,200 years ago (CI: 125 – 3,044), which conflicts with the much later introduction of barley to North America by European migrants. We speculate that this inconsistency reflects the uncertainty of our parameter scaling and note that the confidence interval of our estimates still concur with a European-based introduction to North America a few centuries ago (19).

In conclusion, the demographic history of *P. teres* f. *teres* recovered using ABC and MSMC2 reflects the introduction of its host, barley, in different locations, highlighting the significant role of historic trading in the dispersal of crop pathogens (Figure 6).

**Figure 6:**
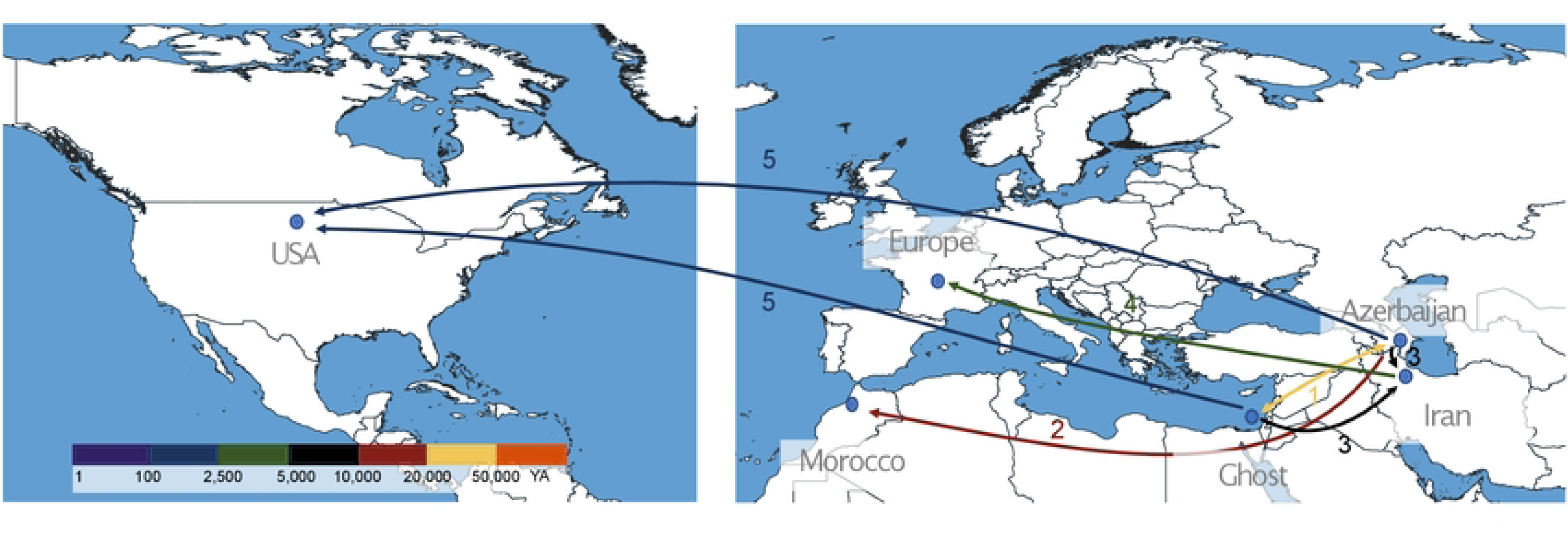
Invasion scenario of *P. teres* f. *teres* based on our population and phylogenomic analyses and the known history of the host. Different colours represent the approximate period when the proposed events occurred. (1) The Fertile Crescent is the most plausible center of origin of the pathogen. (2) Ancestral divergence of the North African population is consistent with an early migration of the pathogen to North Africa, possibly with early barley cultivation by neolithic farmers in North Africa. (3) More recent populations have emerged in Europe and (4) North America.

We used the *P. teres* f. *teres* SNP data to compute overall nucleotide diversity in the pathogen populations. Interestingly, *P. teres* f. *teres* showed higher nucleotide diversity (π) compared to other prominent pathogens such as the wheat pathogen *Zymoseptoria tritici* (65), the wheat powdery mildew pathogen *Blumeria graminis* f.sp. *tritici* (25), and the rice blast fungus *Magnaporthe oryzae* (66). Genetic variation is instrumental in rapid adaptation of pathogens and the high nucleotide diversity in *P. teres* f. *teres* may be an important factor in the successful spread of the pathogen. Differences between species may be explained by different extent of sexual recombination and gene flow and can also be highly impacted by past population bottlenecks. Based on our LD analyses and the distribution of mating type frequencies, we find evidence for frequently occurring sexual recombination in *P. teres* f. *teres* (67). An exception is the North American population that exhibits a higher extent of clonality. The population genetic structure of the North American population is in agreement with a recent founder event of the population, but may also reflect the large-scale monocropping systems in North America that may favour clonal spread of the pathogen over large spatial scales.

We identify six genetic clusters that largely correlate with geographic origin. However, some isolates appear to co-exist and pertain to distinct clusters with little evidence of introgression. This observation may indicate local adaptation and possibly some limits to gene flow, for example between isolates adapted to distinct barley cultivars.

To explore signatures of local adaptation, we used different methods to identify selective sweeps. In total we identify 109 selective sweep regions. We asked how many, and which selective sweeps would be common among the barley-infecting populations; considering that these loci could represent important host specificity loci. Intriguingly, most selective sweeps are population-specific or shared among a small number of populations. Only 23 regions are shared among all *P. teres* f. *teres* populations. Several selective sweeps are shared with the Californian population of *P. teres* occurring on wild grasses. Possibly, genes in these “conserved” sweep regions represent fundamentally important virulence traits.

For the population-specific sweeps, we speculate that the pathogen undergoes strong selection from its local environment, including barley cultivars utilized in different countries. We find that selective sweeps are enriched with predicted effector genes in our regions, which may highlight how host genetics is a main driver of rapid evolution in this pathogen. In addition, as many as 15 out of the 109 selective sweep regions were previously identified as candidate loci in QTL studies aiming to identify virulence determinants in *P. teres* f. *teres* (38,39).

## Conclusion

A growing body of evidence suggests that human activities play a major role in the emergence and dispersal of plant pathogens (25,61,68). Here, we employed population genetic approaches with statistical and simulation tools to unravel the population structure and dispersal history of a major fungal barley pathogen. Our results sediment the conclusion that crop domestication in the Neolithic was accompanied by the emergence of several new plant pathogens; pathogens which co-evolved and spread with their hosts and presently represent some of the most important crop diseases we have. Wild relatives of domesticated plants represent important resources of genetic resistances. Likewise, they may be hosts to “wild populations” of pathogens. Exploring genetic variation in natural plant-pathogen systems holds a large potential for the discovery of new crop resistances as well as pathogen virulence determinants.

## Material and methods

### Genome data

124 *P. teres* whole genomes were sequenced using Illumina technology (PRJNA923641) (69). The sequenced isolates were sampled from barley fields on four different continents. Twenty isolates were obtained from two North Dakota State University experimental fields in Fargo and Langdon, North Dakota, USA. 27 isolates were collected from six locations in Morocco, North Africa. Ten strains were isolated in Iran, and 21 isolates were sampled from five locations in Azerbaijan, South Caucasus. Twenty-one isolates were sampled in Central and Northern Europe and five in Denmark, Europe. Finally, we included a collection of isolates from a wild barley species collected in California, USA. The Californian isolates were included with the purpose of identifying recently diverged genomic features in the barley-infecting populations of *P. teres* f. *teres* (Table S1).

### Read mapping and variant calling

A pipeline was developed to filter and map Illumina reads to a reference genome and extract high-quality single nucleotide polymorphisms (SNPs). In brief, the program Trimomatic version 0.38 (Bolger et al., 2014) was used to filter and trim sequencing adapters, nucleotide bases, based on sequencing quality (PHRED 33), and read length (reads shorter than 30 bp were discarded). Overlapping reads were merged using PEAR version 0.9.11 (Zhang et al., 2014). Burrows-Wheeler Aligner (BWA) version 0.7.17 (72) and Stampy v. 1.0.20 were used (73) to map individual reads to the reference genome of *P. teres* f. *teres* (0-1 *P. teres* f. *teres* genome GCA_000166005.1) obtained from the NCBI (40) (Table S2). Haplotyping and genotyping procedures were performed with the GATK HaplotypeCaller version 4.2.18 (74), providing a final VCF file with the raw SNP calls.

We next conducted filtering of SNPs based on the following criteria: (1) The call quality divided by the depth of sample reads should be larger than 2, (2) the depth per genome should be higher than 8, (3) mapping quality of reads supporting each SNP should be higher than 40, (4) allele-specific rank sum test for mapping qualities of the reference (REF) versus alternative (ALT) reads should be higher than −12.5, (5) allele-specific rank sum test for relative positioning of REF versus ALT allele within reading must be higher than −8. (6) each genome has to have an average read coverage of at least 2. For the application of these filters, GATK VariantFiltration version 4.0.11 was used (74). After applying these hard-filtering criteria, 1,092,635 SNPs were kept, and we further refer to this dataset as the “full high-quality dataset”. We note that additional subsets or filtering steps were added for specific analyses. Most clustering analyses assume that the markers used are independent. Therefore, for these analyses, we filtered the full high-quality dataset based on the linkage disequilibrium (LD) decay patterns considering a distance of at least 3,12 Kbp (distance of r^2^/2 averaged across populations) between SNPs (Table 1). After filtering for LD, a dataset of 465,963 SNPs was retained. We refer to this dataset as the “independent SNP dataset”.

We also generated a dataset of SNPs exclusively located in non-coding, presumably neutrally evolving genome regions. For this, we excluded all SNPs located in predicted gene regions and in 500 bp gene-flanking regions, both upstream and downstream, in a third filtering step based on LD. We obtained the coordinates of the genes from the 0-1 reference annotation file(40). After this filtering step, 160,472 high-quality independent and presumably neutrally evolving biallelic SNPs were kept. We refer to this as the “neutral dataset, which we used to infer the demographic history of the species.

### Population genetic structure

Population genetic structure was inferred using three different approaches: a principal component analysis (PCA), ADMIXTURE version 1.3 (75), and Neighbour-Net analyses (76), each of them based on the “independent SNP dataset.” The PCA was here applied to reveal genetic clustering among the isolates and was created using the R package SNPRelate v. 1.6.4 and visualized with the R package ggplot2 (77). Population structure was further characterized using a maximum likelihood approach implemented in ADMIXTURE. Ten replicate runs for a range of K-values (1-10) were performed. The best K value was determined using the ADMIXTURE software to estimate the cross-validation error(75) (Table S6). Finally, we explored the genetic structure and reticulation patterns among lineages, using the distance-based method for constructing Neighbour-Net networks as implemented in the program Splitstree 4 version 4.15.1 (78). A mantel test was performed to assess the correlation of geographic and genetic distance using the R package ape v. 5.6-2 (79)

### Genetic diversity, neutrality tests and linkage disequilibrium

We further used the “full high-quality dataset” to compute and compare genetic variation among populations. ANGSD v.0.939 (80) was used to estimate the genetic diversity for each population as the nucleotide diversity (π) and the number of segregating sites (Wθ), as well as values of Tajima’s D. The tool PopLDdecay (81) was used with default parameters to estimate the linkage disequilibrium (LD) decay for each genetic cluster. VCFtools v. 0.1.17 (82) was used to calculate the fixation index Fst (83) between the populations. A Kruskal-Wallis test with post-hoc pairwise Wilcoxon was used to identify significant differences (p < 0.05).

### Mating types

*MAT1-1*and *MAT1-2* mating type sequences were obtained from GenBank (accession no. HM121994 and HM122006, respectively) (84). Assemblies were created using SPAdes (44) with default parameters for each isolate. Subsequently, the mating type of each assembly was assessed by blasting the mating type sequences against them. To that end, blastn (85) with default parameters was used.

### Phylogenetic reconstruction

A Maximum likelihood approach was applied to assess phylogenetic relationships between the population isolated from wild barley in California and six others *Pyrenophora* species. Sequences of four DNA loci (ITS, LSU, *tub2*, and *tef1-a*) were extracted from *Alternaria alternata* (SRC11RK2F), *Pyrenophora teres* f. *maculata* (SAMN15340022), *Pyrenophora teres* f. *teres* (SRS084801), *Pyrenophora graminea* (CBS 336.29), *Pyrenophora tritici-repentis* (V0001), and *Pyrenophora seminiperda* (SAMN02981545) from NCBI. Californian isolate CAWB5, showing the highest raw read count among the Californian isolates was selected to represent the group. Subsequently, it was assembled with the software SPAdes (44), following the process described under the “Mating types” section, and the sequences of ITS, LSU, *tub2*, and *tef1-a* were extracted. Consensus sequences of the four individual loci were aligned with MAFFT v 7.490 (86) using default parameters, manually adjusted using Unipro Ugene v. 43.0 (87), and concatenated using SeqKit (88). The concatenated alignment was then subjected to maximum-likelihood (ML) analysis using Iqtree version 2.0.3 (89). The best-fitting substitution model was chosen based on the Bayesian Information Criterion (BIC) using the ModelFinder algorithm implemented in Iqtree version 2.0.3 (90). Moreover, 1000 bootstrap replicates were performed to obtain branch support values using the bootstrap approximation option of Iqtree (91). Further support for the phylogenetic inference was provided by a maximum-parsimony (MP) analysis using MPBoot (92). Similar to ML, MP analysis was performed, including 1000 bootstrap replicates. *Alternaria alternata* (GCF_001642055.1) was selected as the outgroup taxon for both ML and MP analyses. The resulting trees were edited in FigTree 1.4.4.

IQ-TREE polymorphism-aware models (PoMo) (93) were used to reconstruct relationships between *P. teres* f. *teres* populations. For the preparation of the input file, the FastaVCFtoCount.py script provided with the PoMo software was used. Similar to the previous phylogenetic analysis, the best-fitting substitution model was chosen based on the Bayesian Information Criterion (BIC) using the ModelFinder algorithm implemented in Iqtree version 2.0.3. Again, 1000 bootstrap replicates were performed to obtain branch support values using the bootstrap approximation option of Iqtree. The Californian *P. teres* population was used as an outgroup in this analysis.

### Inference of the demographic history of *P. teres* f. *teres* populations

The demographic history of *P. teres* f. *teres* populations was inferred using approximate Bayesian computation (ABC) with a supervised machine learning algorithm implemented in DIYABC-RF version 1.0 (47). Since the ABC framework requires populations without continuous gene flow, the five non-admixed, well-populated clusters revealed by the population structure analyses were used in this analysis. Due to the limited number of individuals in the cluster (only three isolates from Caucasus), this second Caucasus cluster was excluded from the inference. Out of the five remaining clusters, three were entirely consistent with the geographical origin of the populations: North Africa, Middle East, and North America. For the remaining two genetic clusters, the composition of isolates did not reflect on a single geographic location but rather a mixture of isolates from different locations, although these clusters originated primarily from Europe and Azerbaijan, Caucasus (Table 1). Most of the isolates (19/20) of the fourth cluster originated from Europe (France and Denmark). Similarly, the majority (12/18) of the fifth cluster isolates originated from Caucasus. For the inference of the demographic history, we only kept the 19 isolates from Europe, representing the fourth genetic cluster, and the 12 isolates originated from Caucasus, representing the fifth cluster.

Since the records about *P. teres* f. *teres* invasion history are scarce, the phylogenetic analyses obtained with PoMo were incorporated as the starting point to construct hypothetical evolutionary scenarios. First, the three populations (North Africa, Middle East, Caucasus) that were further apart from each other on the tree, indicating ancient split and isolation among these, were selected as a starting point. Three sequential DIYABC-RF analyses were performed as follows: For the first analysis, 49 scenarios were tested, describing cases where (1) either of the single population or (2) an admixture event between two populations gave rise to the other populations (figure S4).

Considering a wider geographical distribution of *P. teres* f. *teres* not covered by our sampling, we also included scenarios that tested an unsampled (referred to as a “ghost”) population as the putative ancestral population. We included scenarios where either a present-day sampled population was derived from a ghost population or the present-day population emerged by admixture from a ghost population with another sampled population. These scenarios were analyzed individually and combined in groups of similar scenarios (47). For the combined groups, scenarios were joined into thirteen groups based on the population that was most ancestral (Table S8): Groups 1,2, and 3 consist of scenarios considering the Caucasus, North Africa, and Middle East population as the origin, respectively. Groups 4,5 and 6 consider North America, Middle East, and Caucasus to have been established through an admixture event of the other two populations, respectively. Group 7 considers that all three populations diverged at the same time. Groups 8 to 13 consider a ghost population to be parental to one of the sampled populations. A detailed description of the scenarios and scenario families can be found in the Supplementary material S1 and Table S7.

In the second analysis, 17 scenarios were considered to assess the emergence and relationship of the European population in relation to the putatively ancestral populations from Caucasus, Middle East, and North America (Figure S5). Group 1 consisted of scenarios where Europe and Caucasus have the same ancestral population (scenarios 1,8,9,16). Group 2 considers Europe and Middle East share a common ancestor (scenarios 2,13). Group 3 considers Europe and North Africa share a common ancestor (scenarios 3,6,14). Group 4 considers that Europe and the “ghost” population share a common ancestor (scenarios 4,7). Group 5 (scenarios 10, 11, 12) consider the European population to be the product of admixture of two other sampled populations. Group 6 considers scenario 17, where the Europe population is the “ghost” population identified in step 1. The best scenario, selected by random forest in analysis two, was used as the base for analysis 3.

In the third analysis, we assessed the relationship of the population originating from the North America with the rest of the population (Figure S6). As many as 26 scenarios were tested. As in the previous steps, group 1 consisted of scenarios where North America and Caucasian have the same ancestral population (scenarios 1,8,9,16). Group 2 considers North America and Middle East to share a common ancestor (scenarios 2,13). Group 3 considers North America and Europe to share a common ancestor (scenarios 3,7,10,23,24). Group 4 considers North America and North Africa share a common ancestor (scenarios 4,8,11,13,25). Group 5 considers the North America and the “ghost” population to share a common ancestor (scenarios 5,9,12,14,15). Group 6, consisting of scenarios 6,7,8,10,11,13 considers the North America population to be the product of admixture of two other sampled populations. Group 7, consisting of scenario 26, considers the North America population as the “ghost” population identified in step 1.

A uniform prior distribution for all effective population sizes in the three analyses ranged from 500 and 5,000,000. The changes in effective population sizes were split into recent changes, where the uniform prior distribution was between 10 and 20,000 years ago, and ancient, where the distribution was between 10 and 1,000,000 years ago. The later distribution was used for all splitting and admixture times with uniform probability. For the scenarios that consider the origin of a population through admixture, the prior distribution of the contribution of each parental population was set to be between 0.01 and 0.99. The random forest analysis for the model choice and the parameter inference was performed with 1000 decision trees and default values of DIYABC-RF. Furthermore, DIYABC-RF uses Hudson’s ms simulator (94) with the “-s” parameter to introduce a fixed number of segregating sites under each scenario. Mutation parameters are, in this case, not needed for the simulations (47). Eventually we have scaled the time values by estimating the current effective population sizes based on Watterson’s theta and a mutation rate of 4,5 × 10^−7^ (25) (Table S9).

MSMC2 v2.1.1 (50) was applied to infer changes in the effective population size through time. MSMC2 uses a backward-in-time algorithm to build back genome lineages. The MSMCtools bamCaller script was used for the preparation of mask files for the low-coverage regions and to “diploidize” the haploid vcf files. After that, the script generate_multihetsep.py included in the MSMC2 software package was used to create the input files for the analysis. Changes in effective population size were inferred using all the isolates available for each population (from 7 for Middle East to 21 for North Africa). Subsequently, the cross-coalescent rate between the populations was estimated using 100 iterations.

### Genome scans for selective sweeps

Three independent approaches were applied to identify signatures of selective sweeps along the *P. teres* f. *teres* genome. Hereby we used the programs SweeD (53), 2), OmegaPlus v. 3.0.3 (54), and 3) RaiSD v 2.9 (55). The analyses were conducted with the full high-quality dataset. SweeD v. 3.0 uses the Site Frequency Spectrum (SFS) patterns of SNPs to estimate a composite likelihood ratio (CLR) test for detecting complete sweeps (Pavlidis et al., 2013). We used SweeD, OmegaPlus, and RaisD individually for each genetic cluster and each of the 12 chromosomes and a grid size equal to the number of SNPs present in each chromosome (28,698 – 77,617 points). OmegaPlus is a scalable implementation of the ω statistic (95) that can be applied to whole-genome data. It uses a maximum likelihood framework and utilizes information on the LD between SNPs. For OmegaPlus, the minimum and maximum window sizes were set to 1 kb and 100 kb, respectively. RaiSD computes the *μ* statistic, a composite evaluation test that scores genomic regions by quantifying changes in the SFS, the levels of LD, and the amount of genetic diversity along the chromosome (55). We used RaiSD with the default window size of 50 kb.

Changes in genetic variation and LD along the genome are also influenced by demography. To account for the effect of demography and determine the significance level of the identified selective sweeps, we simulated 10,000 datasets under the best neutral demographic scenario using the program *ms* (94), to mimic a population evolving under the same conditions as *P. teres* f. *teres*, but without any effect of selection. We then computed the ω and *μ* statistics on this data. Setting a significance threshold for the deviation of the ω and *μ* statistics based on the simulated data sets allowed us to control for the effect of the demographic history of the population on the SFS, LD, and genetic diversity along the genome (53,96). Subsequently, we only kept the selective sweep regions with evidence of selection from at least two methods to control for false positives. Genome-wide maps of the sweep regions were created using Circos v. 0.69-9 (97).

To test if the abundance of effector genes was different for the predicted selective sweep regions compared to the rest of the genome, we used a permutation test (based on a custom script available in GitHub: https://github.com/Jimi92/Population-genomics-Pyrenophora-teres). In brief, the abundance of predicted effectors was counted in regions of the same size and number equal to the predicted sweep regions. As many as 10,000 replicate runs of random resampled region were performed.

### Functional annotation of genes under selection

To predict the potential function of the genes located in the predicted selective sweep regions, we employed eggnog-mapper version 2 (98). To this end, we have used the parodical method for gene prediction with default parameters. For the annotation, the method used the eggnog 5.0 orthology data (99).

Gene ontology (GO) enrichment analysis was performed in shinyGO (100). GO terms were assigned to *P. teres* f. *teres* total genes list in eggnog, which were used as the background list for enrichment analysis. A GO category was considered significantly enriched only when the p-value for that category was < 0.05 after applying FDR correction.

### Co-localization of selective sweep regions, QTLs and predicted effectors

QTLs associated with P teres teres virulence were published in previous works (38,39,101). In addition, effector prediction was performed in a previous work (102)We have compared the candidate selective sweep regions obtained through our analyses to the reported QTL and predicted effector coordinates using BEDTools version 2.27.1 (103)

## Data Availability

Data supporting the findings of this work are available within the paper and its Supplementary Information files. Genome sequences are accessible through Zenodo under (DOI: 10.5281/zenodo.8183372). Custom scripts and workflows are available at https://github.com/Jimi92/Population-genomics-Pyrenophora-teres.

## Acknowledgements

Benoit Barres and Asieh Vasighzadeh kindly provided isolates from France and Iran which were used in this study. The authors are grateful to Idalia Rojas Barrera, Danilo Pereira, Wagner Fagundes and other members of the Environmental Genomics group. This work was supported by the DFG Research Training Group RTG2511.

## List of Supplementary Tables

Table S1: Sampling sites of NFNB.

Table S2: Read count: raw and after filtering for quality, length and PCR duplicates, percentage of reads mapped to the reference genome and mean coverage per isolate across the genome.

Table S3: Pairwise Wilcoxon test to assess significant differences in genetic diversity levels between *P. teres* f. *teres* populations

Table S4: Pairwise Wilcoxon test to assess significant differences in Tajima’s D between *P. teres* f. *teres* populations

Table S5: Proportion of each mating type per population

Table S6: Cross-validation error over 10 replicate runs, the average error per K-value and the standard deviation.

Table S7: Overview of scenario groups used in the three DIYABC-RF analyses.

Table S8: Overview of random forest vote results for each of the three ABC-RF analyses performed for individual scenarios and scenario groups.

Table S9: Estimations of splitting time between *P. teres* f. *teres* populations.

Table S10: Genomic regions that have undergone a recent selective sweep for the Middle Eastern population. The reported regions have been identified by three independent methods (see Methods).

Table S11: Genomic regions that have undergone a recent selective sweep for the North American population. The reported regions have been identified by three independent methods (see Methods).

Table S12: Genomic regions that have undergone a recent selective sweep for North African population. The reported regions have been identified by three independent methods (see Methods).

Table S13: Genomic regions that have undergone a recent selective sweep for the Caucasian population. The reported regions have been identified by three independent methods (see Methods).

Table S14: Genomic regions that have undergone a recent selective sweep for the Californian population. The reported regions have been identified by three independent methods (see Methods).

Table S15: Genomic regions that have undergone a recent selective sweep for the North and Central European population. The reported regions have been identified by three independent methods (see Methods).

Table S16: Pairwise population divergence estimates (Fst) of *P. teres* f. *teres* populations.

## List of Supplementary Figures

Figure S1: Tajima’s D of *P. teres* f. *teres* populations in each geographic region. Kruskal-Wallis test with post-hoc pairwise Wilcoxon was used to identify significant differences (p < 0.05) between the groups(Table S4)

Figure S2: NeighbourNet tree shows the evolutionary relationship between *P.teres* f. *teres, P.teres* f. *maculata* and *P. tritici-repentis*.

Figure S3: Mantel test shows significant corelation between geographic and genetic distance (Number of permutations: 9,999, Pearson corelation coefficient).

Figure S4: Models tested in DIYABC-RF analysis 1

Figure S5: Models tested in DIYABC-RF analysis 2

Figure S6: Models tested in DIYABC-RF analysis 3

Figure S7: GO enriched biological processes and molecular functions. Biological processes and molecular functions identified as enriched in the selective sweep regions. A GO category was considered significantly enriched only when the p-value for that category was < 0.05 after applying FDR correction.

Figure S8: The effect of sample size on the genetic diversity estimations.

